# Brain age prediction of healthy subjects on anatomic MRI with deep learning : going beyond with an “explainable AI” mindset

**DOI:** 10.1101/413302

**Authors:** Paul Herent, Simon Jegou, Gilles Wainrib, Thomas Clozel

## Abstract

**Objectives:** Define a clinically usable preprocessing pipeline for MRI data

Predict brain age using various machine learning and deep learning algorithms

Define Caveat against common machine learning traps

**Data and Methods:** We used 1597 open-access T1 weighted MRI from 24 hospitals.

Preprocessing consisted in applying : N4 bias field correction, registration to MNI152 space, white and grey stripe intensity normalization, skull stripping and brain tissue segmentation

Prediction of brain age was done with growing complexity of data input (histograms, grey matter from segmented MRI, raw data) and models for training (linear models, non linear model such as gradient boosting over decision trees, and 2D and 3D convolutional neural networks).

Work on interpretability consisted in (i) proceeding on basic data visualization like correlations maps between age and voxels value, and generating (ii) weights maps of simpler models, (iii) heatmaps from CNNs model with occlusion method.

**Results:** Processing time seemed feasible in a radiological workflow : 5 min for one 3D T1 MRI.

We found a significant correlation between age and gray matter volume with a correlation r = -0.74. Our best model obtained a mean absolute error of 3.60 years, with fine tuned convolution neural network (CNN) pretrained on ImageNet.

We carefully analyzed and interpreted the center effect.

Our work on interpretability on simpler models permitted to observe heterogeneity of prediction depending on brain regions known for being involved in ageing (grey matter, ventricles). Occlusion method of CNN showed the importance of Insula and deep grey matter (thalami, caudate nuclei) in predictions.

**Conclusions:** Predicting the brain age using deep learning could be a standardized metric usable in daily neuroradiological reports. An explainable algorithm gives more confidence and acceptability for its use in practice. More clinical studies using this new quantitative biomarker in neurological diseases will show how to use it at its best.

**FOREWORD:** *About Owkin:* OWKIN was co-founded in 2016 by Thomas Clozel, MD, a clinical research doctor and former assistant professor in clinical hematology and Gilles Wainrib, PhD, a pioneer in the field of Artificial Intelligence in biology. OWKIN passed the proof-of-concept phase and is now providing its innovative AI algorithms to several of the largest cancer centers and pharmaceutical companies in Europe and in the US. With offices in New York and Paris, we pride ourselves in building a company culture around transparency, collaboration, challenge, optimism and fun.

*Owkin’s team:* Owkin’s team is international, multidisciplinary with incredible talent in machine learning, medicine and business. Our data scientists are among the best in the world, with several Kaggle Masters (top global 100), a DREAM Challenge top performer, and publications in ICML, NIPS and other top scientific journals.

*Tasks repartition:* Idea : Thomas Clozel, Roger Stup, Simon Jegou, Paul Herent Bibliography : Paul Herent, Simon Jegou, Thomas Clozel Data access : Simon Jegou, Paul Herent Data cleaning : Simon Jegou, Paul Herent Data analysis : Simon Jegou, Paul Herent Data preprocessing : Simon Jegou, Paul Herent Data analysis : Simon Jegou, Paul Herent Training of models : Simon Jegou, Paul Herent Work on interpretability : Simon Jegou, Paul Herent Writing : Paul Herent Rereading : Simon Jegou, Thomas Clozel, Julien Savatovsky, Roger Stupp, Olivier Elemento, Kim Gillier Submission to medical congress : Paul Herent, Simon Jegou

Thanks to… Simon Jegou, for your mentoring in Machine Learning, Thomas Clozel and Gilles Wainrib, for your welcome at Owkin, very benevolent, Roger Stupp, for your support and re-reading, Julien Savatovsky, for your support and re-reading, Valentin Ame and Sylvain Toldo, for your help on the beautiful figures and the design of the related <u>blogpost</u>, All the Owkin team members, for the great team work we did (and hope we’ll do) between Paris and New York : Anna Huyghues Despointes, Anna I. Bondarenko, Pierre Courtiol, Derek T. Russell-Kraft, Cedric Whitney, Meriem Sefta, Vincent Lepage, Adrian Gonzalez, Maxime HE,Paul Jehanno, Raphaël Léger, Alicia Simion, Eric Tramel, Mikhail Zaslavskiy, Pierre Manceron, Chloé Simpson, Paul Mabillot, Valentin Amé, Mathieu Galtier, Camille Marini, Sylvain Toldo, Charlie Saillard, Olivier Dehaene, Olivier Moindrot, Pascal Roux, for your support, your help, your advices, Axelle, for your patience and support.

## INTRODUCTION

### Limits of nowadays radiology ?

A part of work of radiologist today can be described as a perceptual and a cognitive task : as a perceptual task, the physicians has to recognize features considered as normal or not. As a cognitive task, he has to describe relevant features in a written radiological report, and finally conclude, providing to the patient and his physicians what’s his global interpretation : a set of hypotheses in a diagnosis exam, an evolution of disease compared to a previous exam…and guidelines for the following medical treatment.

Some feature interpretations are very well known, and consensually classified and correlated to a known risk. For example in Breast Cancer medical imaging, American College of Radiologist (ACR) defines the Bi-Rads classification (Breast Imaging Reporting and Data System), (Monticciolo et al. 2017), which aims to classify pattern of radiologic features in 6 classes (from 0 to 5) corresponding to 6 differents probabilities of malignancy. So, The Bi-Rads provides an evidence based decision tool for the physician, indexed on a probabilistic risk of malignancy (practice a follow up exam, make a biopsy…). However, having reliable results is not an easy task.

#### Argument of Replication Crisis in Medical Research

In 2016, a survey published in *Nature* sheds light the problem of reproducibility in Science (BAKER, s. d.). In this survey, 1576 researchers were asked about replication crisis via a brief questionnaire. More than 70% of them have tried and failed to reproduce another scientist’s experiments, and more than half have failed to reproduce their own experiments. More focused on Medicine Research, same pattern of responses occurred : respondents reported between 55 and 75 % of failure in reproducing an experiment. Based on this fact, it seems difficult to call “Evidence-based medicine” a not reproducible evidence based medicine, despite the existence of rating methods for quality of evidence, such as for example GRADE guidelines (Balshem et al. 2011).

#### Argument of medical errors

Errors in medicine represent a significative prevalence (rate of missed, incorrect or delayed diagnoses estimated to be as high as 10%-15%) (Bruno, Radiographics, 2015).

More specifically In radiology, the reported retrospective error rate among radiologic examinations was 30% (Lee et al. 2013), inducing high cost for health systems : errors has been estimated to more than 38$ billion annually in the US.

In the sus-cited review (Bruno, Radiographics, 2015), the three most frequent causes of errors related concerned missed finding (42% of related errors), satisfaction of search (a finding is missed because of failure to continue to search for additional abnormality after the first abnormality was found (22% of related errors)), and faulty reasoning (ie misclassification (9% of related errors)). Among the causes of errors, we can mention implication of fatigue (Reiner, 2012, J digit imaging), lack of ergonomic tool, organisation issues because of increasing of volume of medical images demand in medical practice… This errors could explain the high inter and intra observer variability observed in radiologic interpretation (Muenzel et al. 2012), (Suzuki et al. 2010).

### What is desirable for the future of radiology ?

#### Radiology has the potential to go beyond human visual perception

Relevant information is present in imaging data which are humanly difficult to quantify. The radiological data can be implanted in models including other data (clinical data, genomic data, histological data). A new discipline, called Radiomics, has been created to answer to this challenges. It can be defined as a “high-throughput mining of quantitative image features from standard-of-care medical imaging that enables data to be extracted and applied within clinical-decision support systems to improve diagnostic, prognostic, and predictive accuracy” (for a review, see (Lambin et al. 2017)). Some radiomics tools have already shown to be efficient. One good example is texture analysis. It’s humanly difficult to quantify a degree of heterogeneity in a tumor for instance, whereas texture analysis algorithms can. Moreover, some evidences showed that’s method can reflect at a tissue scale a genomic mutation. This could have a great impact to accelerate prediction medicine and avoid cost of tissue sequencing. Some recent evidence showed the interest of this technique in predicting EGFR mutation in lung adenocarcinoma CT scan (Sacconi et al. 2017)), or IDH1 mutation in low grade glioma MRI (Zhou et al. 2017).

#### A strong paradigm change with rising of Artificial Intelligence in Radiomics filed

Some confusions can be made in the definitions : today what is called “artificial intelligence” in medicine concerns mostly supervised machine learning tasks. Deep learning is a subtype of machine learning techniques, called “deep” because of the digital architecture using an important number of layers of artificial neurons (LeCun, Bengio, et Hinton 2015). Majority of deep learning papers are using convolutional neural networks, secondary of its success observed in Computer Vision challenge (Russakovsky et al. 2014), where this techniques outperformed standard computer vision algorithms.

Since, some proofs of concept has been published in computer vision tasks relative to medicine in histology (for a review see (Litjens et al. 2016), for a recent paper published by our team, see (Courtiol et al. 2018)), dermatology (Esteva et al. 2017), ophthalmology (Gulshan et al. 2016), radiotherapy (for a review, see (Meyer et al. 2018))…

Field of applications of deep learning in radiology is potentially very large and could revolutionize each step of medical imaging pipeline : image reconstruction (Zhu et al. 2018), image segmentation, medical interpretation (caracterisation of lesion, prognostic prediction…). For reviews, see (Yasaka et al. 2018), (Hosny et al. 2018).

As mentioned by the white paper of Canadian society of radiology focused on artificial intelligence (Tang et al. 2018), the combination of improved availability of large dataset combined with increasing computing power permit machine learning techniques such as deep learning to rapidly moving from an experimental phase to an implementation in medicine. Moreover, managing clinically machine learning projects for clinical practice need a combination of expertise between data scientist and data expert. This work is the result of this collaboration.

### A black box ? The need of explainable Artificial Intelligence

To be implementable in clinical practice, Algorithms need to be understood.

There is a well known trade off in the Machine learning community between Performance in Interpretability. A simpler model is more interpretable, but less accurate. A more complex one can produce high level performance (like deep learning), but is criticized as a black box. Moreover, AI systems are very sensitive to bias, and its prediction can be the result of capturing confonds instead of answering to the asked question.

To overcome this lack of common sense and for a better acceptability in clinical practice, a model has to be as accurate and explainable as possible. The real challenge is here.

For a review on different models usable in medicine, see (Holzinger et al. 2017).

### Why Brain Age Prediction ?

#### Brain Ageing physiology

Physiology of brain ageing is classically studied in two manners : structural and functional ways. In a structural way, brain ageing studies focused on studying ageing features of grey matter, white matter, Cerebro-Spinal Fluid (CSF), and vessels. For grey matter, morphometric techniques such as voxel-based cortical thickness (VBCT), and voxel based morphometry (VBM) permitted to quantify cortical atrophy in different brain regions (Hutton et al. 2009). For example in (Good 2001), with VBM technique, local areas of accelerated loss were observed bilaterally in the insula, superior parietal gyri, central sulci, and cingulate sulci, whereas no effect of ageing was observed in amygdala, hippocampi and entorhinal cortex, showing a pattern of heterogeneous atrophy in brain.

Focusing on white matter, Leukoaraiosis showed to be a good predictor of physiological ageing, reflecting the presence of small vessel disease. Some correlations were found between leukoaraiosis and cognitive decline (Grueter et Schulz 2012). Its physiopathology is multifactorial, with role of inflammation, blood brain barrier disruption, genetic factors, ischemic factors (for a review see (Lin et al. 2017)).

Brain Ageing was also studied with functional MRI. Functional connectivity permitted to identify brain regions associated with cognitive reserve (defined by discrepancy between clinical symptoms and effects of aging in Alzheimer Disease). For instance regions such as medial temporal regions and anterior or posterior cingulate cortex were associated with neural reserve, and neural compensation (ie alternative cognitive strategy when a cognitive dysfunction occurs) in frontal regions and dorsal attention network (for a review, see (Anthony et Lin 2017)).

#### Brain Ageing in radiology

Today, in radiological routine, physiological ageing assessment consists in mostly 4 tasks, such as (i) categorical assessment of atrophy (eg : medial temporal lobe atrophy score (MTA), aiming to distinguish normal to pathologic atrophy (Gaillard s. d.)), (ii) assessment of ventricles dilatation (the task consists in identifying atrophy to hydrocephalus), (iii) assessment of White Matter Hyperintensities (WMH) on MRI T2 sequences (this tasks consists in firstly identifying the leukoaraiosis pattern vs other WHM patterns (for eg : inflammatory, such as in multiple sclerosis), and secondly grading level of leukoaraiosis with Fazekas scale (Fazekas et al. 1987)), and (iv) detect, count, and assess spatial distribution of cerebral microbleeds (CMBs), in order to distinct normal ageing in the brain to the main differential diagnostics : high cardiovascular risk (deep spatial repartition CMBs are associated with small vessel disease due to hypertension) versus Amyloid angiopathy (superficial, lobar distribution of CMBs) ; for a recent review see (Haller et al. 2018).

Schematically, 3 MRI sequences are used for this 4 tasks : T1 weighted MRI for grey matter and CSF, T2 weighted (FLAIR sequence) for white matter (and CSF), T2* or SW for CMBs assessments. In this paper, we focused only on T1 weighted images, as a preliminary step. Having a biomarker of physiological ageing could be a clinically relevant in radiological routine. For instance, Brain age prediction was assessed in Multiple Sclerosis, showing difference between healthy people and patients (Raffel et al. 2017) : the brain of patient with MS had predicted age 10 years “older” with this biomarker).

### Brain Age prediction : a potential biomarker in diseases

Brain age prediction from MRI consists in applying a classical machine learning process : try to predict Y (the brain age) from X (the MRI data) with a model computing a function f(X) for predicting Y.

A recently published paper focused precisely on this topic (Cole et al. 2017), providing to us a state-of-the-art result for this task : Error prediction was 4.65 years (Mean Absolute Error, MAE) with 3D convolutional neural networks (CNNs) trained on raw T1 weighted brain MRI (ie minimal preprocessing) with a dataset of 2001 subjects.

Brain Age Prediction with machine learning has been studied as a biomarker of accelerated ageing in some diseases like schizophrenia (Koutsouleris et al. 2014), multiple sclerosis (Raffel et al. 2017), traumatic brain injury (Cole et al. 2015), mild cognitive impairment (Gaser et al. 2013) and Alzheimer disease (Franke et Gaser 2012).

Having a brain-predicted age indicative of an older-appearing brain was associated with known biomarker of ageing : motricity (weaker grip strength), breath (poorer lung function), cognition (slower walking speed, lower fluid intelligence), marker of chronic stress (higher allostatic load) and increased mortality risk. (Cole et al. 2017),

Brain Ageing is an heterogeneous process. Notably, different studies focused on grey matter morphometry showed accelerated pattern of ageing in specific brain regions, such as Insula (Good et al. 2001) or deep grey matter such as thalami and caudate nucleus (Fama et Sullivan 2015). Our work on interpretability with an occlusion method from (Zeiler et Fergus 2014) permitted to observe a similar pattern.

Some papers focused on the interpretability of CNN in related work such as in an Alzheimer disease classification task on structural MRI (Yang, Rangarajan, et Ranka 2018) showed that cortical regions (particularly in the hippocampus), and ventricle were important for prediction. However, to our knowledge, no work on interpretability with similar method was used on healthy subjects.

### Objectives of this study

- Define the key preliminary steps of a Machine Learning project : access to data, data cleaning, data analysis.
- Define a clinically usable preprocessing pipeline for brain age prediction.
- Reproduce previous results (prediction of brain age with 3D CNN) and try to outperform them.
- Compare the performances of 3D CNN with simpler models commonly used by the Machine learning community (linear models, non linear models, 2D CNN with transfer learning)
- Compare the performances of CNN with different data inputs (ie different features from Brain images, growing in dimensionality and complexity (histogram of intensities, grey matter segmented, raw data with minimal preprocessing, effect of skull stripping on prediction)
- Reveal some Caveat against common machine learning traps.
- Work on interpretability of models : from simple models to CNN.

## METHOD

### Getting data

We downloaded two open-access dataset and collected 1597 T1 weighted MRI from healthy subjects : dataset 1 from <u>IXI</u>) (563 subjects) and dataset 2 from <u>Functional</u> <u>Connectome Project</u> (1054 subjects).

Precisions on sequences parameters are described in the links above. On data set 1, MRI were performed on 1.5 tesla and 3 tesla. On dataset 2, majority of MRI were performed on 3 tesla MRI.

### Data cleaning

First step when acquiring one or more dataset consisted in cleaning data in order to check some basics stuff : remove missing data, fixing errors in ID from tabular data and corresponding file.

### Data analysis

Figure 1 is a summary of the data-analysis we made. figure 1A shows the discrepancies of number of patients depending on the center, and relative size of both dataset. Figure 1B shows the heterogeneity of age distribution.

**Figure 1:**
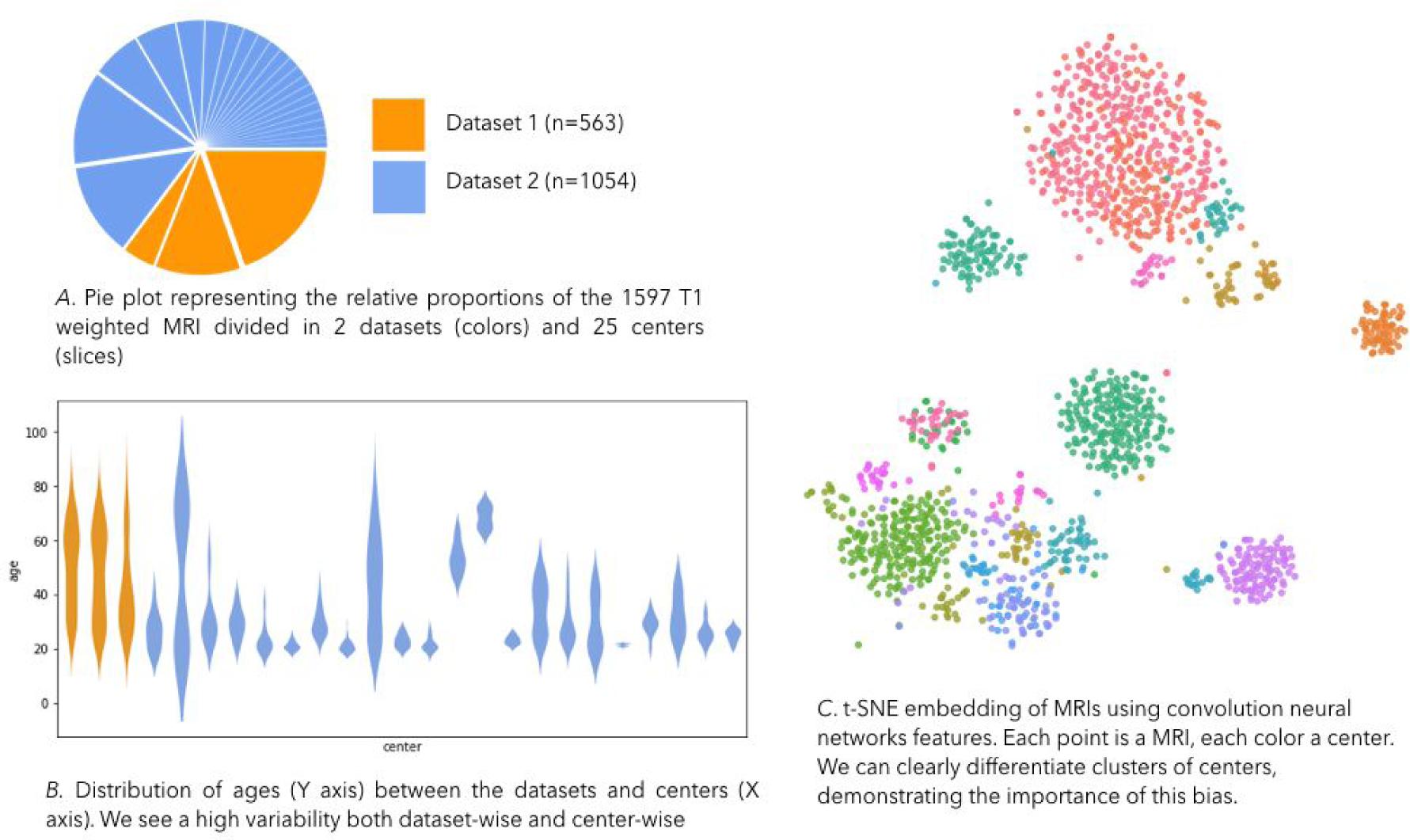
Dataset description.

On figure 1C, we performed a non supervised machine learning task, called t-SNE (t-stochastic neighbor) embedding, in order to identify clusters present in the dataset. This method permitted to identify a strong bias relative to the center were acquired the MRI, such as the number of subjects by centers (hospitals), distribution of age, gender, but mainly due to image acquisition : type of MRI used, size of Field of view (fov) for brain acquisition, defacing method employed to anonymise the dataset, variations in Signal to noise ratio due to tuning of parameters used for T1 weighted MRI sequence, magnetic field strength…

This step permit to show the interest of standardization : we don’t want that our machine learning models capture the biases relative to the center for predicting age (for example, there are centers with outliers (very old subject), if our model identify features relative to this center (ex : contrast specific to the MRI employed), it will help it to predict extreme age, independently of the feature known to be present with brain ageing (atrophy, leukoaraiosis…). For that, multiple strategies can be adopted for controlling this confond factor : augment the size of the dataset (when its possible) with more centers ; data augmentation (ie showing to the model the same images in training phase, but in a different way (doing some basic operations such as rotations, translations of images, etc)) ; method of cross validation for splitting data between train set and test set (Cf Method section).

### Preprocessing

A first key point is that before training the model, an important pre-processing step remained necessary. This took for us the majority of time investigated for this project. We proposed here a minimal preprocessing pipeline, in order to control common biases seen in raw data. This tools are open-source and quite easy to implement. A schematic summary presents this aspects in *figure 2.*

**Figure 2:**
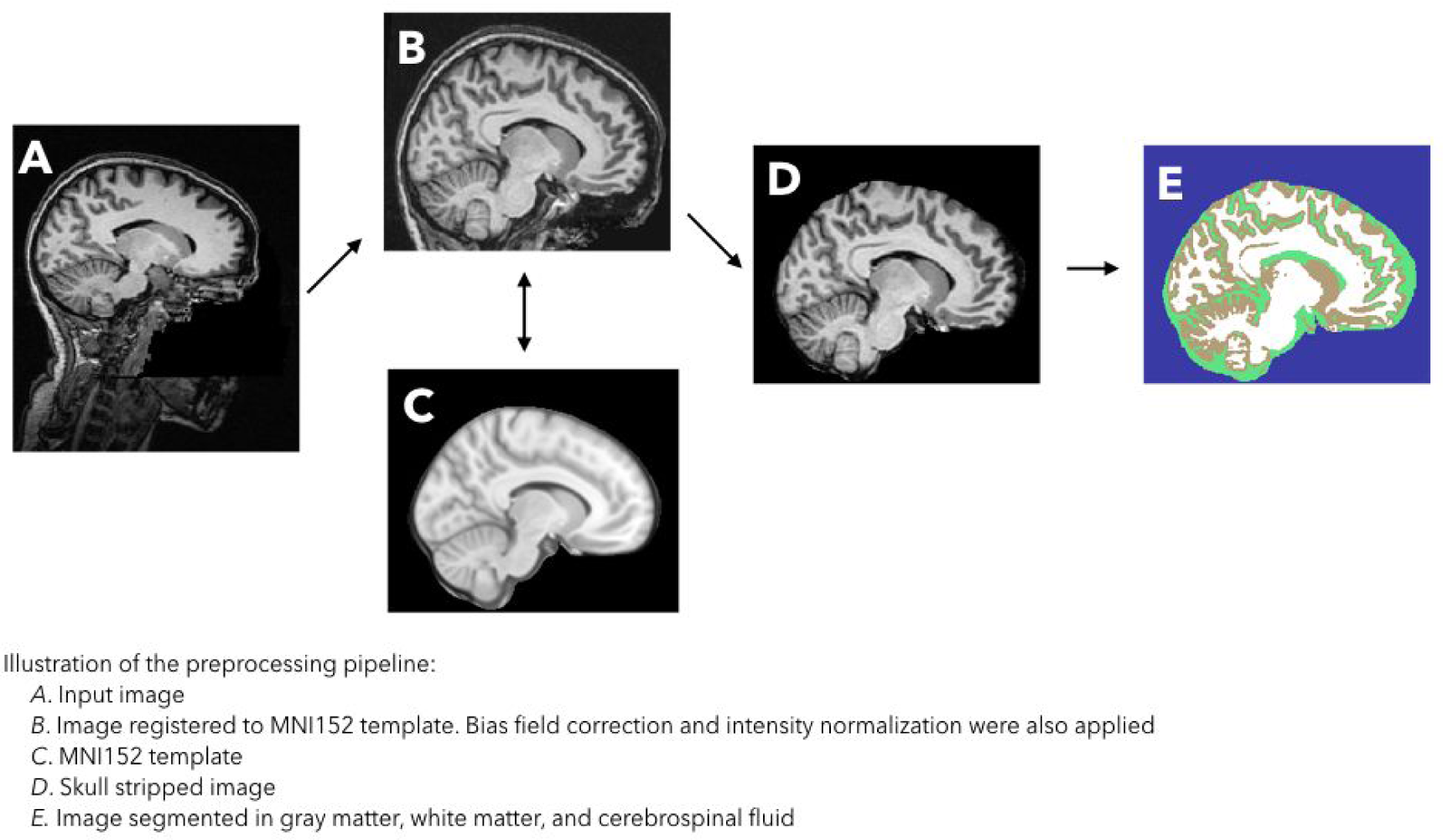
Schematic view of Preprocessing pipeline.

#### Cropping

In *Figure 2.A*, we can observe raw data. From this sagittal view, we can see that defacing was made in order to anonymise subjects for open-access. The neck is seen in this image, but was not always visible, depending on the FOV employed. A standardized cropping centered around the brain was necessary.

#### Co-registration

In *Figure 2. B,* in order to automate this cropping, we employed co-registration tools. We used the MNI152 template for that (template is shown in figure 2.C). MNI (Montreal National Institute) template is a brain atlas which is the average of 152 normal MRI scans that have been matched to the MNI305 using a 9 parameter affine transform (Evans et al. 1993).

3D Spatial transformations are of two types : linear (rigid transformation and affine), and non linear (most used : diffeomorphic transformation). Co-registering in MNI template permits to normalize the relative size of each brain, without losing information about the relative proportion of atrophy (as we saw previously, atrophy mechanisms are not homogeneous in different brain regions). This was also a way to control gender bias (women brains tends to be smaller than man (Ruigrok et al. 2014).

#### Bias field correction

Bias field correction consisted in correcting voxel intensities due to the presence of a low frequency intensity nonuniformity (for details, see (Tustison et al. 2010) (cf *figure 3*). This step was performed using Ants (Advanced Normalization Tools) (http://stnava.github.io/ANTs/).

**Figure 3:**
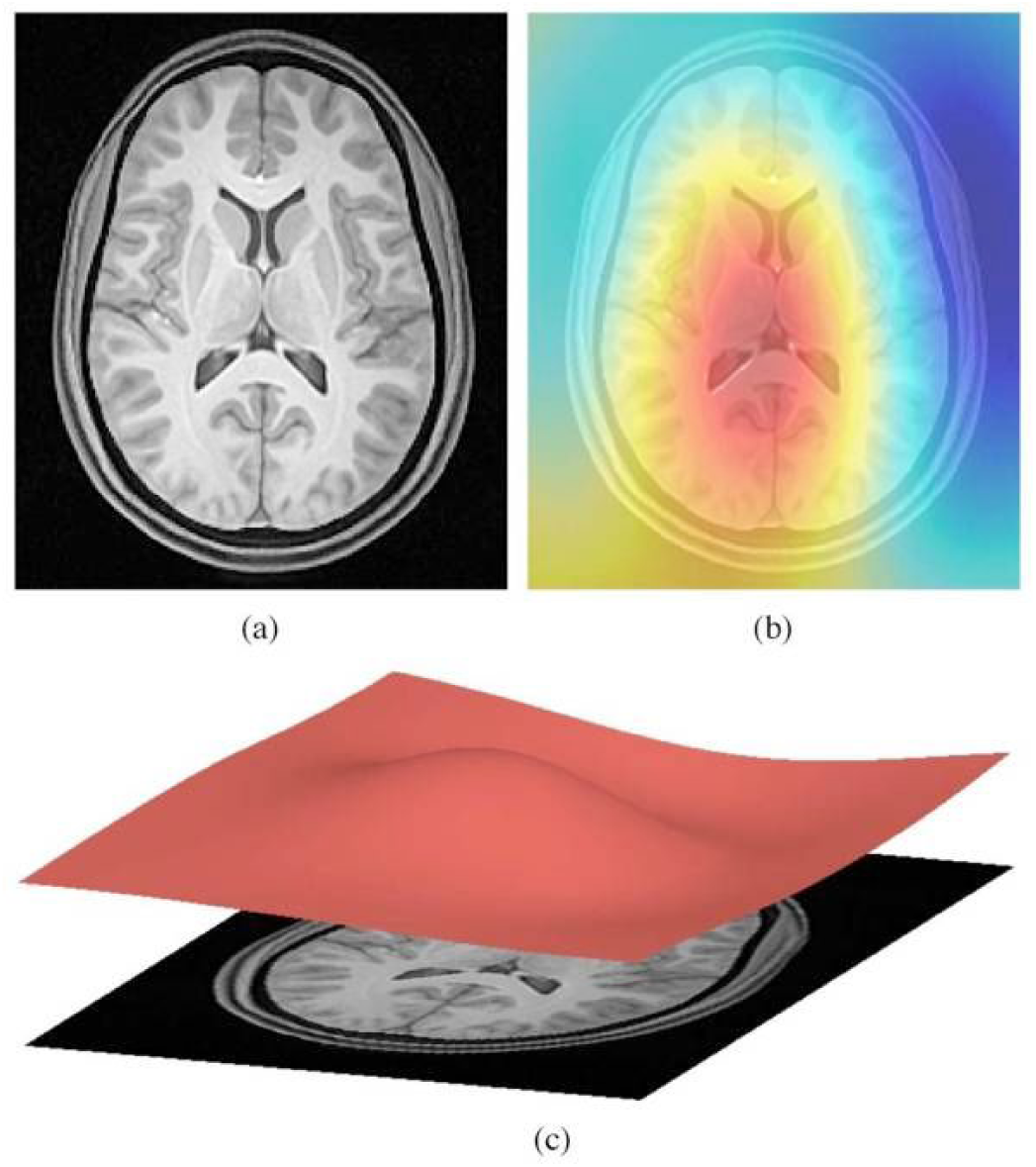
Heat map of inhomogeneity in pixel intensities before N4 bias correction. Image from (Tustison et al. 2010)

#### Skull stripping

Skull stripping consisted in removing the non brain tissue from image (cf *Figure 2.D*). This step was made in order to assess the effect of the presence of non brain tissue (muscle, skin, eyes, bones) on age prediction.

We downloaded the mask of MNI. Then for each image we multiplied the image per this mask. Thus all values in the image with a 0 associated in the mask were removed. Histograms of intensities reflects indirectly the Volume of grey matter. in fact, the most grey matter you have, the highest peak you have in the grey matter range of intensities. Histograms can be used as a basic feature extractor. We reduced brain images to this representation and trained the models in order to constitute a “Baseline” method for age prediction.

#### Intensity normalization

In MRI, there is no absolute value for pixel intensities, contrary to CT scan (were pixel intensities are correlated to the density of tissue, in Hounsfield Unit). Relative contrast is assessed between tissue : In brain, on T1 weighted sequences, grey matter is darker than white matter (cf *figure 4*), in other words the value of grey matter voxel is lower than white matter. The consequence of that is huge variability between values depending on the hospital where images were acquired. A simple way to counter this effect is to normalize intensities centered to a maximum intensity in Brain. First (cf *figure 5*) we centered around 1 the value of pixels on a local maximum corresponding to white matter, which is hyperintense in T1 weighted image (understand “hyperintense” as more intense than grey matter). For that we detected white matter peak in image (with a maximum detection algorithm) and we divided the value of pixel intensities by the value of this peak.

**Figure 4.**
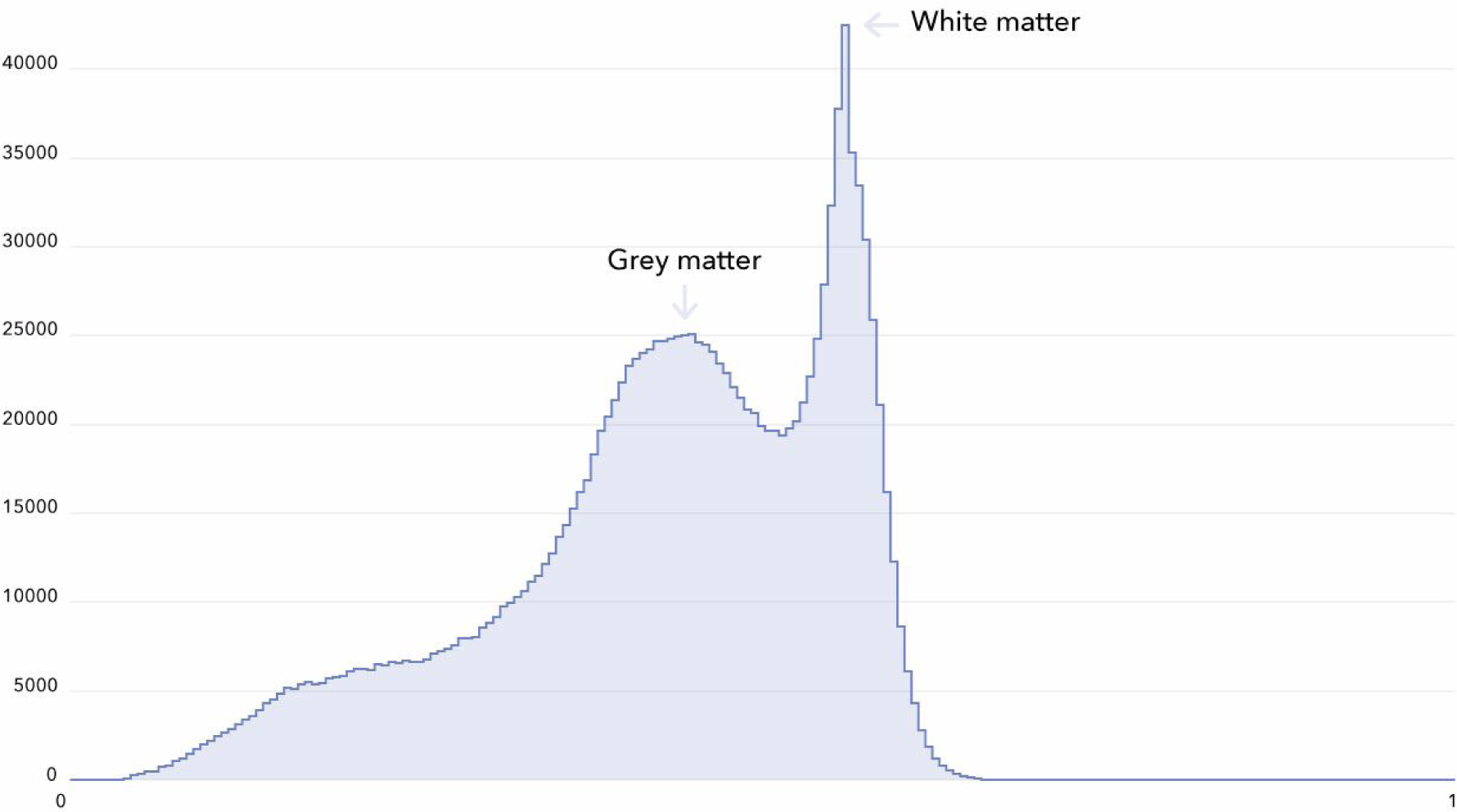
Histogram of intensities in MRI data from the whole MRI : Intensity normalization was made around to the White matter peak of intensities, seen at the right of the image.

**Figure 5.**
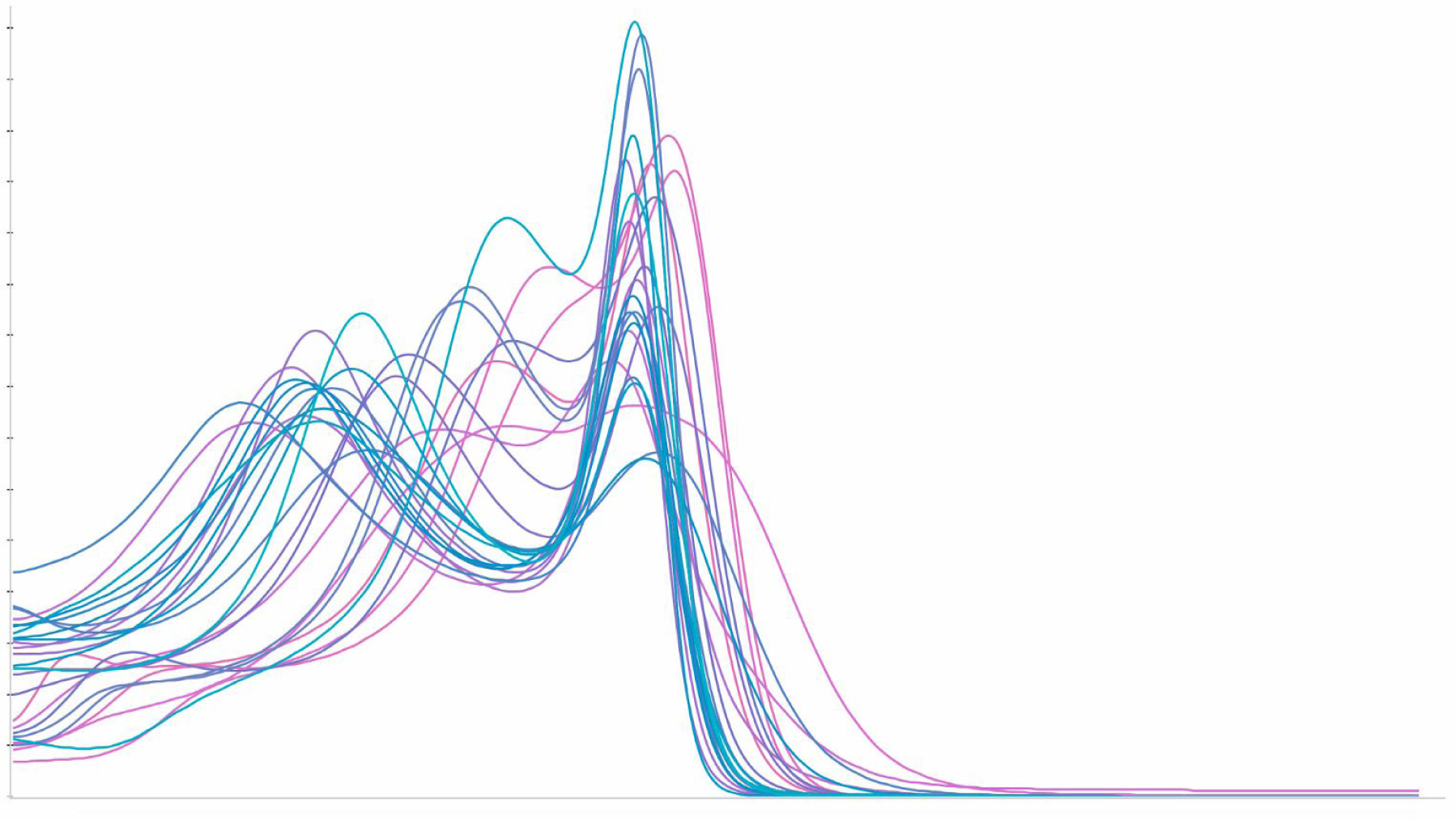
Figure showing the histogram (v1) plots after white matter intensity normalization.

In a second time, we performed an improvement of intensity normalization, which permitted to have a constant point for the local maximum corresponding to grey matter (cf *figure 6*). For that obtained the grey and white matter peak from the segmentation masks. We normalized on white matter peak (same method than before), and secondly interpolated the pixel value between 0 and 1, in order to have the grey peak at value of 0.75. We then assessed this more advanced normalization on performances.

**Figure 6:**
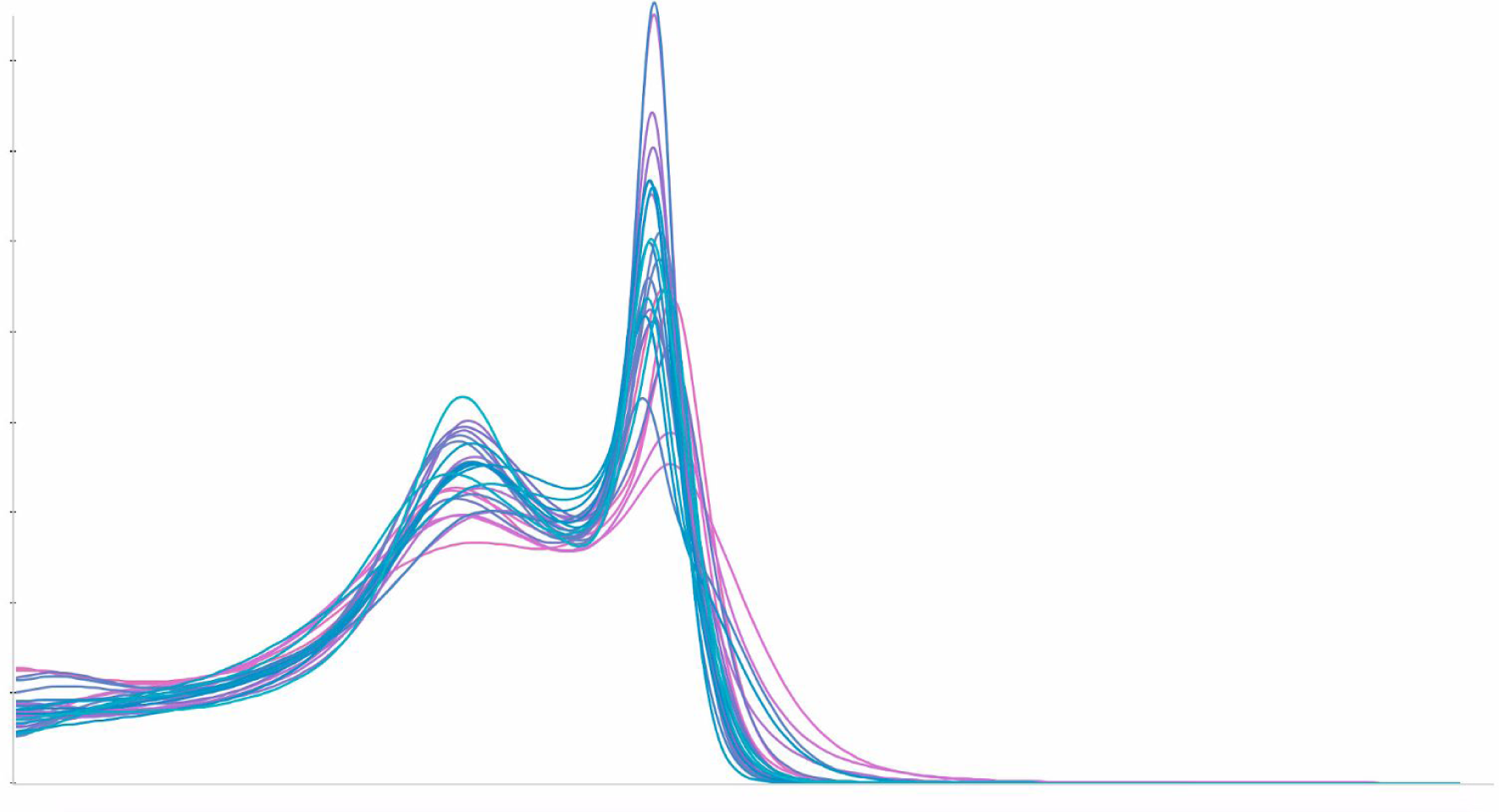
Histogram (v2) of intensities after normalization on Grey and White matter peak intensities.

##### Segmentation of brain Tissue

As you can see in *Figure 2.E*, this final step consisted in separating grey matter to white matter and CSF.

Similarly to histograms, segmentation of different tissues is a way to reduce the dimensionality of data, ie to reduce the complexity. From continuous signal values between grey matter, white matter and CSF, we reduce the pixel intensities in categorical values : 0 for background, 1 for CSF, 2 for grey matter, 3 for white matter.

With this method, we kept the spatial correlations between the different elements of brain tissue, but we lost subtils variations of signal in each type of tissue. We first used Ants library but we were not satisfied of the result. We finally used the FSL toolbox for this step (Zhang, Brady, et Smith 2001).

This step permitted to show correlation between age and grey matter volume (cf *figure 7*). Grey matter is known to decrease with age (Good et al. 2001). We calculated Pearson correlations between age and volume of grey matter from segmentation (ie the sum of voxels corresponding to grey matter). Same calculus were made with CSF and white matter. As expected, a strong negative correlation was observed between age (Y axis) and grey matter volume (X axis) (pearson = -0.74, p<0.05). Correlations were weaker between age and CSF (pearson = 0.40) and White matter segmentation masks (pearson = 0.13).

**Figure 7:**
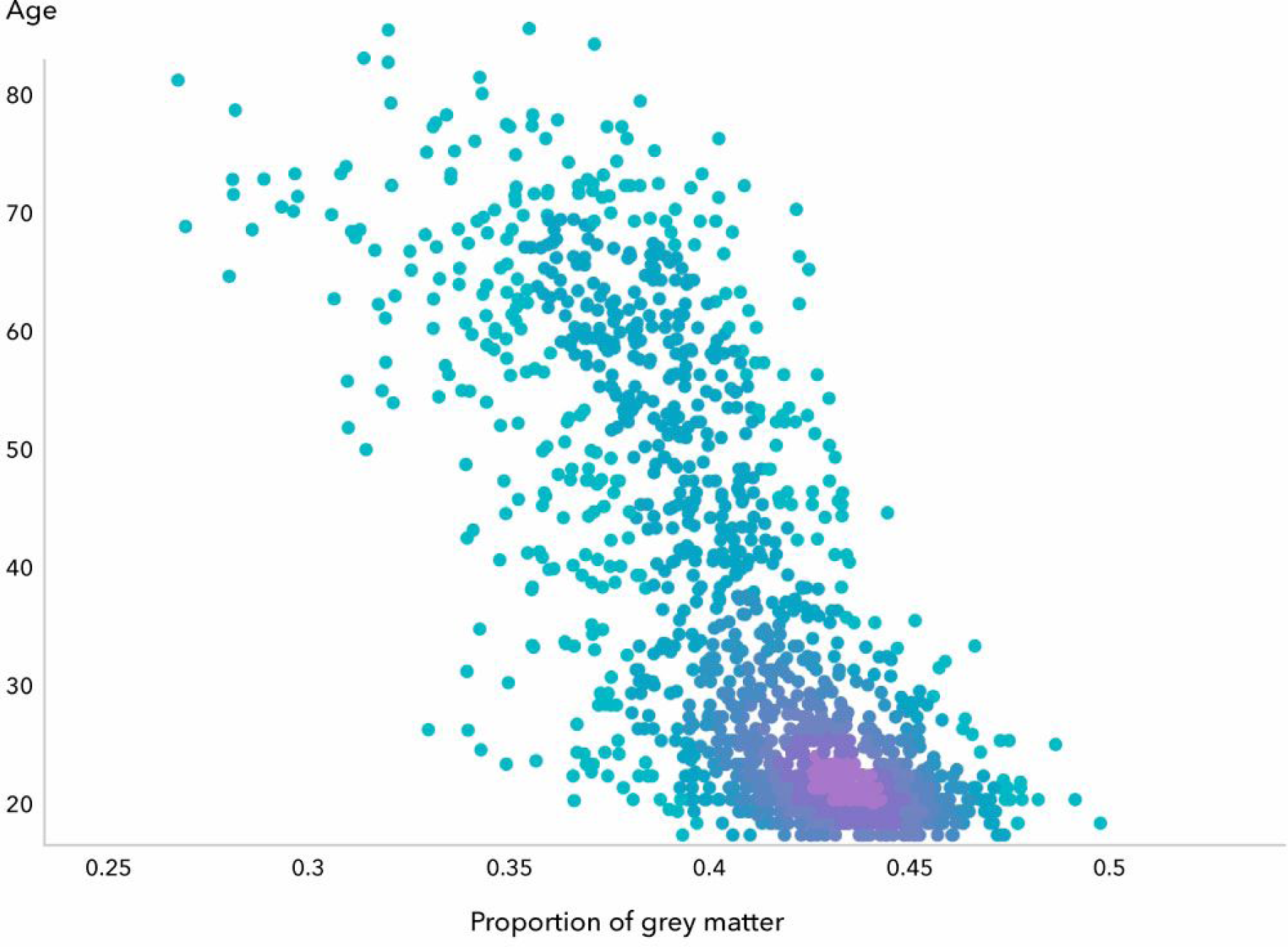
correlation between age and grey matter volume from segmentation masks.

### Machine learning models : Predict Y from X, with different architectures, and the interest of Cross Validation

#### Feature engineering

At this step we have a clean dataset, and pre-processed data in different ways. We have reduced the dimensionality of data and extracted features in different manners in order to have different data inputs : histograms of intensities from MRI, skull stripped segmented masks, skull stripped raw data. In order to assess the effect of skull stripping on predictions, whe have also trained models with non skull stripped raw data.

#### The interest of baseline methods

We trained linear models (linear regression, Ridge Regression) with different data inputs : histograms of intensities from the whole MRI, grey matter mask, and raw data. We also trained models on non linear model : Random forest with Gradient Boosting : CatBoost (Prokhorenkova et al. 2017).

Boosting is a meta-algorithm (ie an algorithm which is applied on other algorithm) based on the idea of gradually aggregating numbers of simple algorithms, called weak learners, to get a final strong learner (Freund et Schapire 1997). More specifically, each weak learner is optimized to minimize the error on the training data using the sum of the previous weak learners as an additional input. While deep learning methods have proved to be extremely efficient on “structured” data (images, sound, text, time-series etc.), boosting methods using decision trees (in fact CART trees) as weak learners remain the non-linear default algorithm used by machine learning practitioners for any other type of data.

#### Complex non linear model : Convolutional neural networks (CNNs)

Deep Learning is a family of non-linear models called neural networks, that share in common a fully differentiable and layer-structured architecture. Deep learning methods have been recently replaced in the spotlight for their impressive performances in computer vision (Krizhevsky, Sutskever, et Hinton 2017), speech recognition (Graves, Mohamed, et Hinton 2013), natural language processing (Bahdanau, Cho, et Bengio 2014), and reinforcement learning (Silver et al. 2016), and these successes have been attributed to three main factors : the availability of larger datasets, of more computing power, and of more sophisticated algorithms.

From a mathematical point of view, one of the key ingredient that makes neural networks so efficient has been the ability to integrate differentiable operations well suited to the structure of the data. In Computer Vision in particular, the use of Convolutional Neural Networks constitute a promising tool. In this paper, we used some of the latest tools from deep learning, such as residual connections (He et al. 2015) and batch normalization (Ioffe et Szegedy 2015), into fully connected neural networks. We trained 2D CNN models separately with in inputs : Segmentation mask of Grey matter and raw data with skull stripping.

All models were trained on single GPU (NVidia Titan Xp or Geforce GTX 1080), and written in Keras.

#### 2D CNN

We used architecture with 2D CNN. We trained models from scratch, and also tried transfer learning with pretrained network (Resnet50), and finally tried data augmentation. The architecture of the 2D CNN trained from scratch contained 4 repeated blocks of a (3x3) convolutional layer, a 2D Batch Normalisation layer, a rectified linear unit (Relu), a 2D Average pooling Layer. The number of feature maps was set to 16 in the first block, and was doubled after each pooling layer to infer a sufficiently rich representation of the brain. The final age prediction was obtained by using a fully connected layer, which mapped the output of the last block to a single output value.

In each application, the network weights were trained by minimizing the Mean Absolute Error (MAE) using a RMS propagation and a Nesterov momentum optimizer (Nadam). The principle of transfer learning (using a pre trained neural network like Resnet 50) is described as follow : it is possible to transfer to a new task the way an algorithm extracted features from millions of image to a new dataset (in this case : Imagenet, a dataset which was trained with 1.4 million of images). In this paper, we removed the 2 last layers from Imagenet, and add two convolutional layers, in order to fine tune them with the specificity of our dataset.

Data augmentation consisted in applying operations on histograms (sharpness, contrast and brightness), an rotations (+/- 5 °), width and height shift of +/- 5 pixels), zooming (factor of 0.01). The interest of data augmentation is to show artificially more data to the algorithm. The more data you have, the best performance you can theoretically obtain (ie a better generalization). We notably focused on augment the variety of histogram values in order to limit the bias due to this heterogeneity coming mostly from image acquisitions.

#### 3D CNN

Data input had the same shape (182x218x182 voxels). 3D CNN followed architecture from (Cole et al. 2017) in order to reproduce the results (for a schematic representation, see *figure 8*). It contained 5 blocks of (3x3x3) convolutional layer, a ReLU, a (3x3x3) convolutional layer, a 3D batch normalization layer, a ReLu and finally a (2x2x2) max-pooling layer. The number of feature maps was set to eight in the first block, and doubled after each pooling layer. The final age prediction was obtained using a fully connected layer, which mapped the output of the last block to a single output value.

**Figure 8:**
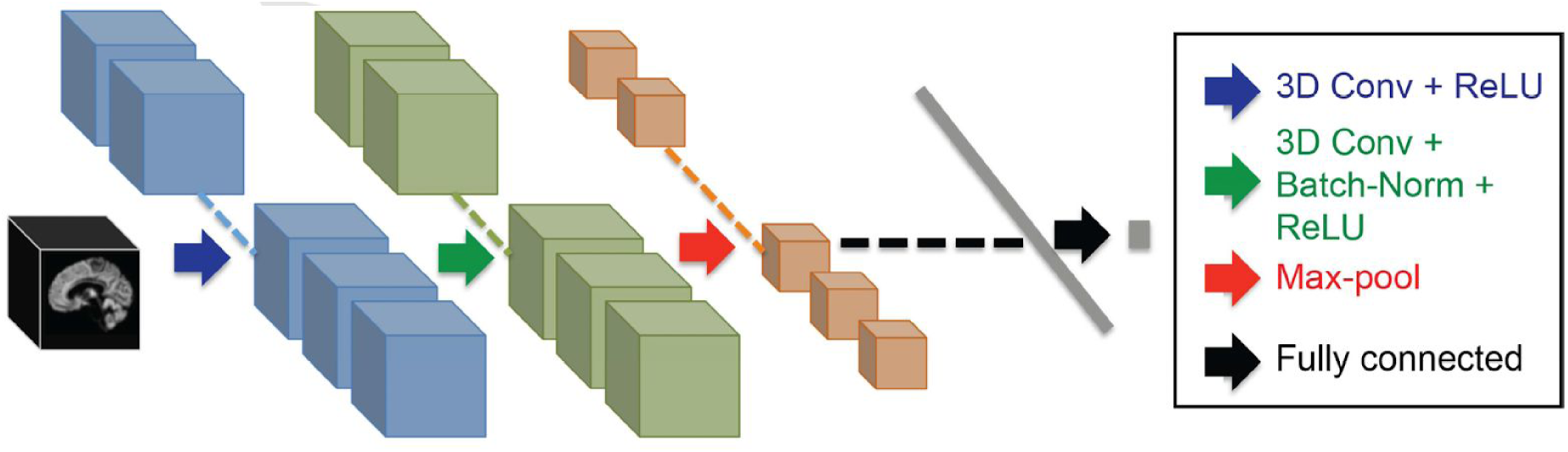
Schematic representation of 3D CNN from (Cole et al. 2017)

In each application, the network weights were trained by minimizing the MAE using different optimizers : Stochastic Gradient descent (SGD) ; Root Mean Square propagation and gradient momentum (ADAM) and a variant of ADAM (ADAMAX). We kept the default tuning parameters proposed by Keras for each experiment.

##### Interest of using different cross validation methods

The random split consisted in a K-fold cross validation with 5 folds (k = 5). 4 folds are used for training the model (80 % of the dataset), and the last one was dedicated to testing. We also wanted to control the center effect on prediction. For that, we splitted the data set in 5 fold, defined by their center (ie data from a given hospital couldn’t be both in training and test set).

#### Work on Interpretability

Interpretability of models if a key domain for validity, reliability and acceptability of algorithms in medical care. One of the challenges in deep learning results in finding tools which permit to make them more interpretable.

In this work we focused on the interpretability of simple models. We first studied the voxels which were correlated with age on different segmentation masks). Secondly we plotted heat map in order visualize the regions of interest for prediction made with a simple linear model (ridge regression). Finally, our work on the interpretability of CNN consisted in covering up images in order to reveal the strongest features selected by the model, based on the method of (Zeiler et Fergus 2014).

## RESULTS

### Predictions made on Histograms

**Table 1.** Brain age predictions made on histograms of images. (Mean Absolute Error ²/- Standard Deviation).

| Algorithm | Cross validation method |  |
| --- | --- | --- |
|  | Random Split | Hospital Split |
| Predicting Median age on the dataset | 13.73 +/- 13.49 | 15.12 +/- 11.87 |
| Median age per hospital | 8.66 +/- 9.66 | X |
| Histogram v1 - Linear Regression | 9.08 +/- 11.19 | 14.22 +/- 9.65 |
| Histogram v1 - Gradient Boosting | 5.74 +/- 5.73 | 11.52 +/- 8.67 |
| Histogram v2 - Linear Regression | 7.35 +/- 6.41 | 9.51 +/- 7.35 |
| Histogram v2 - Gradient Boosting | 5.86 +/- 5.94 | 8.67 +/- 7.39 |

The median age was 36 years. We calculated Mean Standard Error (MSE) from the median age of subjects (13.73 year in random split) in order to compare the predictions. In machine learning terms, standard error is equivalent than predicting the mean age of the training set (ie, a constant value).

We can observe that linear regression is less accurate than non linear method (CatBoost Regression here, a random forest regression with gradient boosting). Moreover, the cross validation method has a strong influence on results. Performance are better using a random split, because this method doesn’t take into account the center effect, which act as a bias on performance.

We can observe that more advanced normalization (giving a constant pixel value to grey matter and white matter in histograms) improved performances both for random and center split. Moreover, the differences of performance between random split vs center split were smaller with this method. It can be explained by the fact that this method is a way to control the center effect (notably center specific contrast between grey matter and white matter).

### Prediction on Grey matter segmented mask

In *Table 2*, we show the predictions made with grey matter segmentation masks as inputs of models. We can see that predictions are more accurate than those made with histograms. The performances are equivalent with simple models, and no gain in performance was observed with non linear method such as Gradient Boosting.

**Table 2.** Brain age prediction made on segmented mask of grey matter (Mean Absolute ²/- Standard Deviation).

| Algorithm | Cross validation method |  |
| --- | --- | --- |
|  | Random Split | Hospital Split |
| Histogram V2 - Gradient boosting | 5.86 +/- 5.94 | 8.67 +/- 7.39 |
| Segmented MRI - Linear regression | 4.70 +/- 3.84 | 6.17 +/- 4.75 |
| Segmented MRI - Gradient Boosting | 4.92 +/- 4.38 | 7.09 +/- 5.50 |

With cross validation made by Hospitals, performances are closer between random vs center split (eg : 4.70 years vs 6.17 for Linear regression), suggesting a lower center effect with this method. We could infer that by the fact that segmentation mask removed bias present in signal (exemple : specific signal to noise ratio inherent to the different MRI machines).

### Prediction on Skull stripped raw data with 2D Convolutional Neural Networks

In *Table 3*, we can see results obtained with models trained on skull stripped raw data (by “raw” we want to say we applied minimal preprocessing described in method section : normalization on grey and white matter, bias field correction, co-registration in MNI space, and skull stripping). We can observe that 2D CNN trained from scratch is less accurate than a pre trained one. Data augmentation improved performance both on random and center split. The best result was observed with pre-trained 2D CNN on random split (MAE : 3.60 years), and outperformed state-of-the-art results. However this gain in performance was not observed on center split (MAE : 6.58 years).

**Table 3.** Brain age prediction made on skull stripped raw data with 2D CNN (Mean Absolute Error ²/- Standard Deviation).

| Algorithm | Cross validation method |  |
| --- | --- | --- |
|  | Random Split | Hospital Split |
| Histogram V2 - Gradient boosting | 5.86 +/- 5.94 | 8.67 +/- 7.39 |
| Segmented MRI - Linear regression | 4.70 +/- 3.84 | 6.17 +/- 4.75 |
| 2D CNN trained from scratch | 4.57 +/- 4.72 | 6.94 +/- 5.76 |
| 2D CNN + data augmentation | 4.29 +/- 4.47 | 6.14 +/- 5.47 |
| Pre-trained 2D CNN + DA | 3.60 +/- 3.67 | 6.57 +/- 5.11 |
| Average of CNN + DA and Linear regression | 3.92 +/- 3.55 | 5.50 +/- 4.30 |

### Prediction on Skull stripped raw data with 3D Convolutional Neural Networks

**Table 3.** Brain age prediction made on skull stripped raw data with 3D CNN (Mean Absolute Error ²/- Standard Deviation).

| Lab | Data set<br>(number of<br>subjects) | Skull<br>Stripping | Data<br>Aug-<br>ment-<br>ation | Optimization<br>Method | Cross<br>Validation<br>Method |
| --- | --- | --- | --- | --- | --- |
|  |  |  |  |  | Random Split |
| J Coles,<br>Neuroi-<br>mage,<br>2017 | 2001 | No | Yes | SGD | 4.65 |
| Owkin<br>Lab | 1597 | Yes | No | SGD | 7.10 |
|  |  | Yes | No | ADAM | 5.02 |
|  |  | Yes | No | ADAMAX | 4.78 |

With 3D CNN trained from scratch on skull stripped raw data, we obtained similar results from the state-of-the-art performance (4.78 years vs 4.65 years in the original paper), without data augmentation, with a smaller dataset and using a different optimizer (ADAMAX, a derived version of Adam optimizer, instead of Stochastic Gradient Descent (SGD).

#### Correlations maps on segmented masks

In *figure 9,* we studied the local correlations between segmentations mask voxels and Brain Age. On the left side, only voxels segmented as grey matter are represented, On the right side, only CSF. Blue voxels represent negative correlations, and red voxels positive ones. As expected, cortical area were negatively correlated with age on grey matter segmentation mask (continuous arrow), and positively correlated with subarachnoid space and Ventricles in CSF segmentation mask (“circle” dotted arrow), compatible with the physiology of ageing (atrophy and secondary ventricle dilatation). However, positive correlations were found on grey matter segmentation mask (“square” dotted arrow). This was due to a wrong segmentation of grey matter. Leukoaraiosis is a “grey matter mimics” on T1 sequences (ie in low signal on T1, as grey matter). The segmentation algorithm interpreted this feature as grey matter.

**Figure 9:**
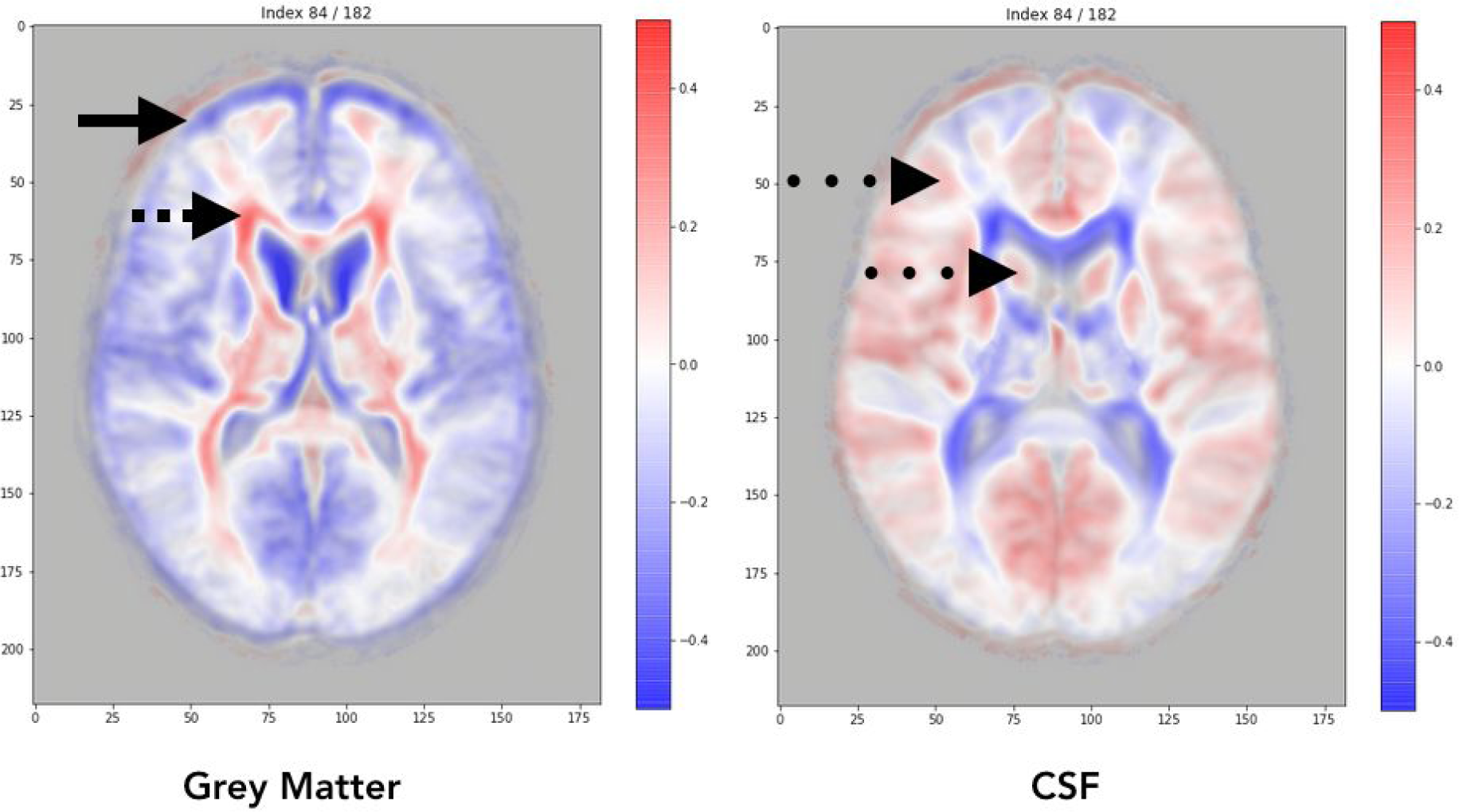
correlations maps obtained on segmentation mask.

In *figure 10,* Red areas represent zones where gray matter was the most correlated with the age according to a ridge regression. The most correlated areas seemed to be the Grey matter and the Ventricles (anterior horn of lateral ventricles and third ventricles). This observation is compatible with the physiology of ageing, combining atrophy associated with secondary and progressive dilatation of ventricles.

**Figure 10.**
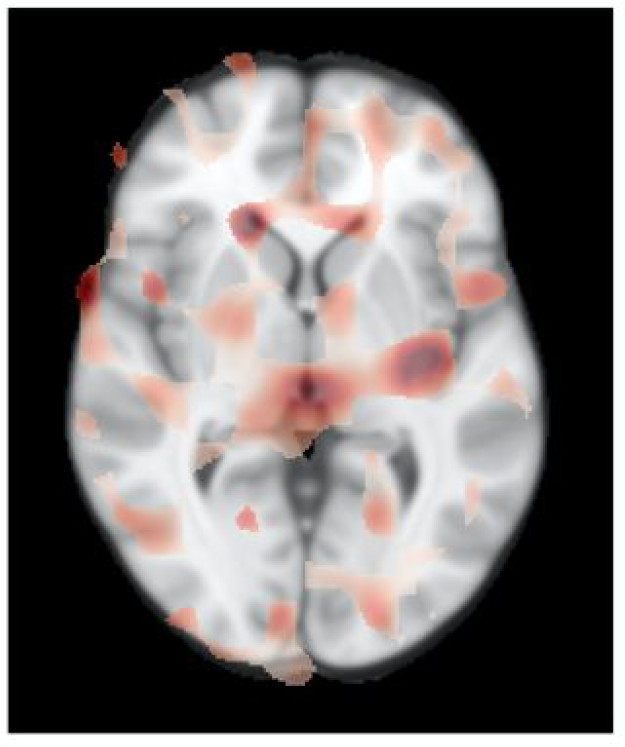
Weights maps from a ridge regression model. Red areas represent zones where grey matter is the most correlated with the age according to a ridge regression.

#### Interpretability of CNN

In *figure 11*, we occluded with small square regions of images and observed their incidence on prediction on pretrained 2D CNN. For a given hidden square, a given prediction error was computed. The more pink the square, the higher the prediction error. On *Left, we* applied this technique on the youngest subjects of the dataset (less than 30 years old). On *Right*, on the oldest (more than 60). We can observe the importance of insula for age prediction. It is consistent with results from (Good et al, 2001), where Brain ageing was described as an heterogeneous process, with some regions known to have “accelerate” ageing, such as in insula.Moreover, deep gray matter (thalami, and internal capsule) was associated with higher prediction errors when occluded, consistent with the results from (Fama et Sullivan 2015).

**Figure 11:**
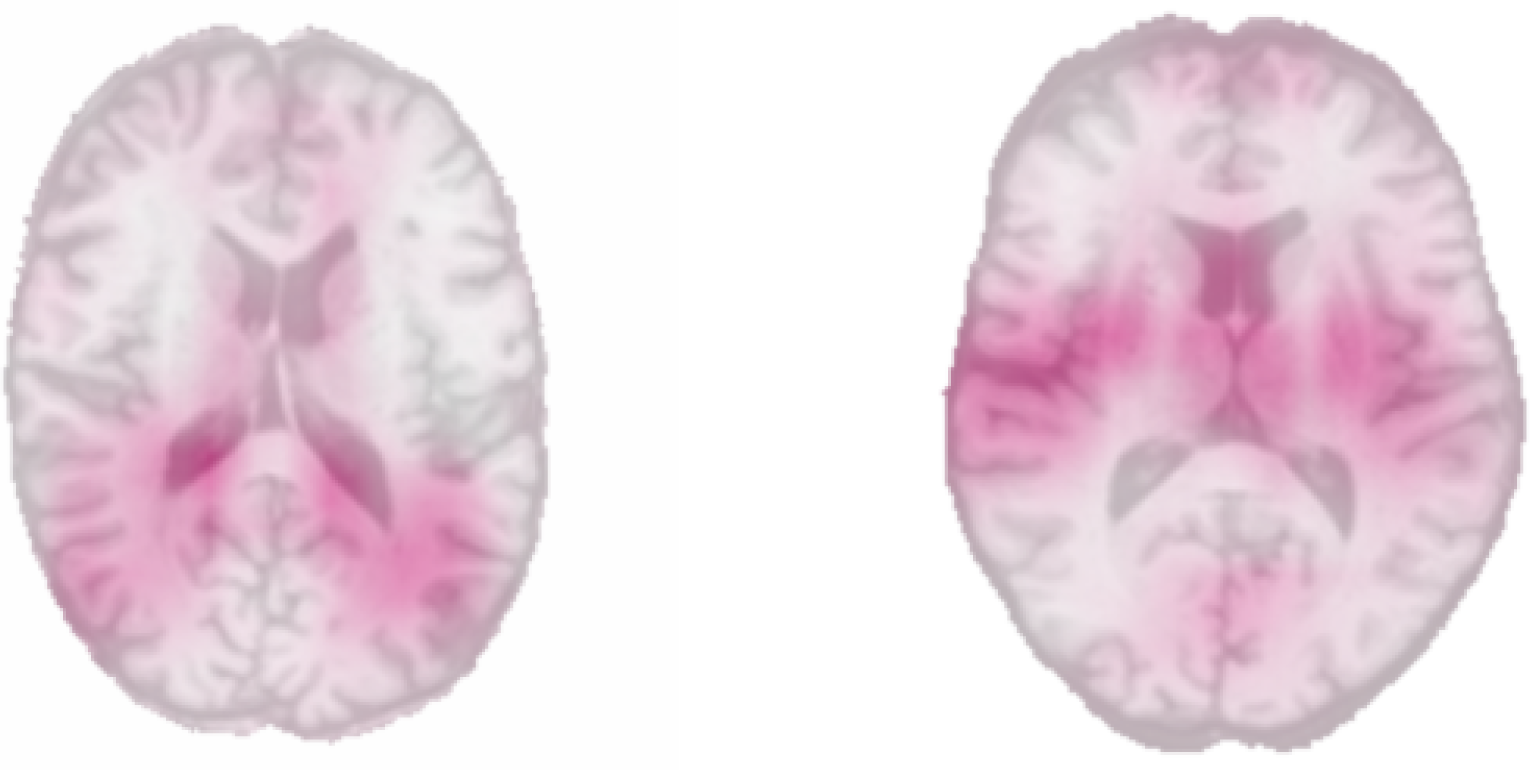
Interpretability of 2D CNN : ”Occlusion maps” with method from (Zeiler et Fergus 2014). On the Left, youngest subjects (less than 30) ; on the right, older ones (more than 60).

In *Figure 12*, on *Left* : we see in this coronal view the « projection » of the error predicted on each sagittal slices (ie each column of pixel in the image is associated with its mean prediction from the corresponding sagittal slice). Prediction was harder on the edges of the brain where less informative data was available.

**Figure 12:**
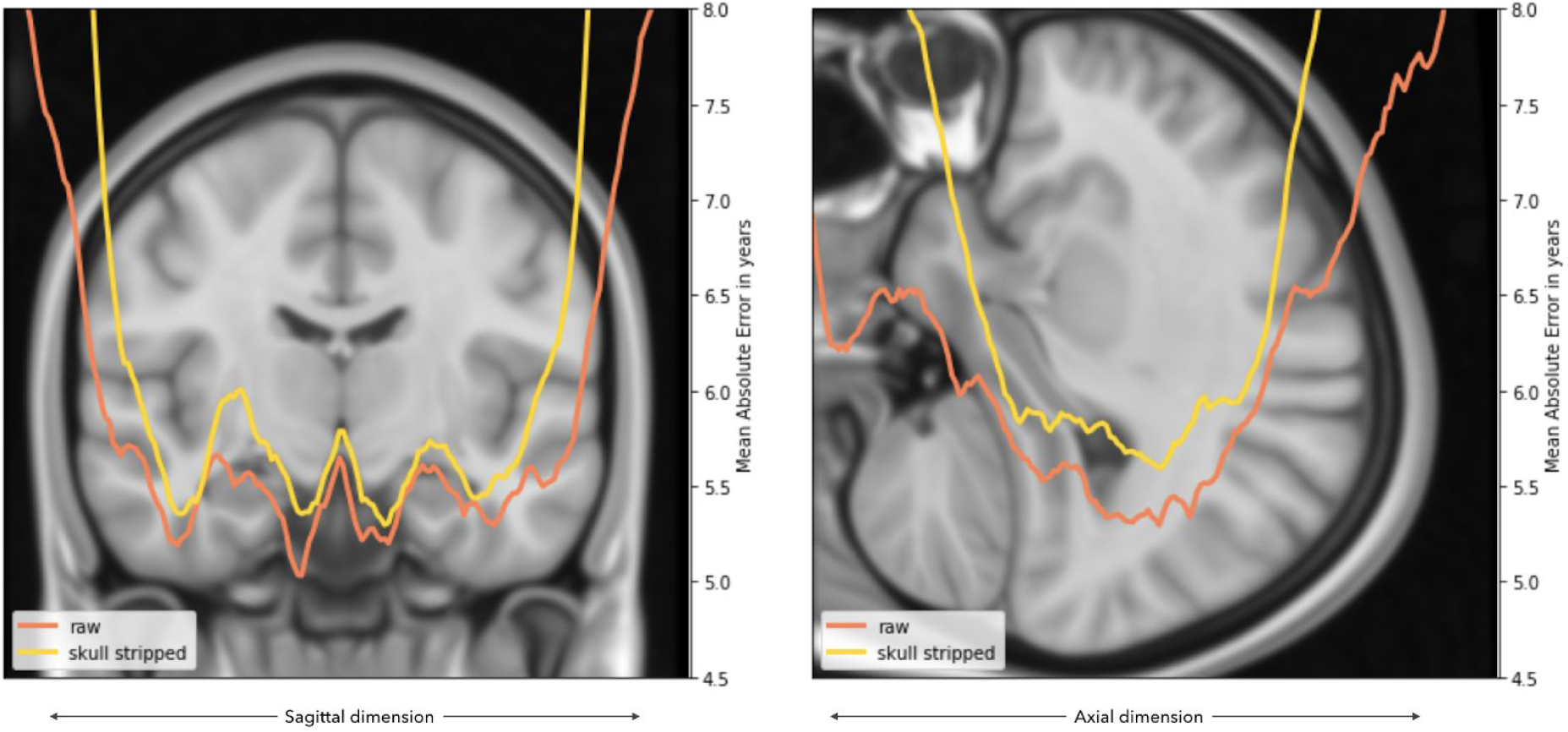
Effect of skull stripping on predictions. We first extract features with Resnet on 2D slices on sagittal or axial plan, on skull stripped data and raw data. We used this features as inputs in a logistic regression model to predict age. We mapped the value of MAE from skull stripped data (yellow curve) and raw data (orange curves) on the corresponding slice in orthogonal plan.

On *Right* : while the error was increasing at the bottom of the brain for the skull stripped curve, it stayed quite low for the raw curve, showing that the model was using features unrelated to the brain (skull, skin…). Thus, all results reported on raw data are unreliable as they predict the age of the head rather than the age of the brain.

## DISCUSSION

### About the results

The tasks consisting in cleaning the data and building an efficient preprocessing pipeline is long and took majority of time. From this preprocessing pipeline we could assess different levels of representation of MRI : from Histograms and segmentation mask of grey matter to raw data. Even the simplest way of representation (histogram of intensities) keeped information about brain ageing, and permitted to obtain results much better than random prediction (best performances with linear Regression) : MAE of 5.86 years vs 13.73 years with random prediction. More accurate normalization was a way to limit the loss of performance on center split.

Linear models were outperformed by non linear models in brain age prediction on histograms, but this tendency was not observed with Grey matter segmentation masks. Simple and non linear models are efficient and get similar results compared to more complex architectures like CNN (Gradient boosting : MAE of 4.70 on grey matter segmentation mask, vs 3.60 years with pre-trained 2D CNN on raw data). CNN outperformed simple non linear models. Pre trained 2D CNN was the best model we trained (MAE : 3.60 years) but was still sensitive to the center effect (MAE : 6.57 years with hospital split). This huge loss of performance was surprising for us. In comparison the 2D CNN trained from scratch was more “resilient” to the center effect (MAE : 4.29 years with random split vs 6.14 years with center split). We observed that the convergence of loss was obtained faster with 2D pretrained CNN. We can hypothesize that the “too fast” convergence resulted in overfitting and bad generalization, resulting a loss of performance on an independent dataset.

3D CNN reproduced similar performance to the ground truth (our best model : MAE of 4.78 years vs 4.65 years on original paper) with less data (1597 vs 2001), but was less performant than 2D CNN in our study. This can be explained by the higher computational cost of using this model. Relevant information was probably sufficient on 2D slices. Interest of using 3D CNN probably depends to the problem to solve. For instance, better performance was observed on a cerebral microbleeds detection task with 3D CNN (Dou et al. 2016), because of the need to use 3D information for differentiating microbleeds from “mimics” (eg : a vessel) on 2D slices.

About the Interest of using skull stripped data :

The best results of original paper (Cole et al. 2017) was obtained using segmented grey matter (which require preprocessing steps of skull stripping and grey matter segmentation) as an input of 3D CNN (MAE : 4.16 years, vs 4.65 on raw data). Author claimed that the use of raw data could permit to avoid long preprocessing steps, and make this kind of algorithm usable in a radiological workflow without preprocessing. In practice preprocessing can be long (hours of computing), depending on tools used and available computational power.

However with efficient computational power, time of processing pipeline can fall down significantly (5 min with our computational power).

The interest of practicing skull stripping remains still unclear, and has to be studied in pathologic cases. On one hand, we can consider that the prediction of age from raw data (ie brain and non brain tissue) constitute a bias : we predicted the “head age” instead of brain age. It could be non relevant for assessing neurologic disease where alteration of tissue is observed only on the brain. On the other hand, the difference in predictions between non skull stripped and skull stripped data are very similar (difference of MAE : less than 0.5 year, cf *figure 12*), and probably constitute an acceptable difference in clinical practice. Moreover, it’s not impossible that the non brain tissue is informative for better predictions in neurologic disease. A specific “mismatch” between pathologic tissue (brain) and healthy tissue (bone, skin, eyes…) could help for defining clinically relevant biomarker of ageing, taking into account heterogeneity of ageing between organs, for instance. This suppose to consider a given disease as an “accelerate ageing” process however.

### About the MRI sequence used in this study

We can reveal some limits of brain age prediction on T1 sequences : (i) the use of unimodal MRI (only T1 weighted MRI was studied here, whereas in radiological routine, a classical minimum Brain MRI protocol contains T1 weighted MRI sequence, FLAIR, DWI, T2*, and TOF) ; (ii) No specific assessment of Leukoaraiosis was done (however this data was captured by grey matter segmentation, cf *figure 9*) ; (iv) Predictions were made only on structural MRI ; (v) the heterogeneity of MRI sequences parameters and the differences of magnetic field strength (1.5 and 3 tesla mostly).

### About interpretability of models

We highlighted the importance of interpretability of models, which permitted to compare our results with known biological pattern of ageing. For less complex models, we highlighted the importance of ventricle dilation and regions were grey matter is situated in the brain. Moreover, these techniques permit to show in a more understandable way the heterogeneity of brain ageing. Moreover, occlusion maps on CNN models revealed the importance of insula and deep grey matter as a relevant regions for predictions. This results are consistent with the known process of accelerated ageing in this regions.

### Brain Age as a neuroradiological biomarker ?

A study using MRIs from 36891 subjects (Kaufmann et al. 2018) showed the interest of predicting age in brain disorders, in particular in pathologies like dementia, mild cognitive impairment or schizophrenia, where a “gap” was observed between the distributions of predictions, compared to healthy subjects.

In another paper (Becker, Klein, et Wachinger 2018), the authors proposed a derived metric from MAE, which aims to quantify the uncertainty of prediction error (covariance). This metric permitted better classification accuracy than the MAE for a classification task such as normal vs Mild Cognitive Impairment (AUC of 0.81 with covariance, 0.68 with MAE), Alzheimer disease (AUC of 0.77 with covariance, 0.68 with MAE), Autism (AUC of 0.60 with covariance, 0.53 with MAE). However this results were not efficient enough to constitute a clinically relevant diagnosis tool (a classical “good” diagnosis test is fixed for AUC around 95%), and is probably due to the unimodality of MRI data (a T1 sequence is a way for studying structural information in brain).

Some authors proposed Multimodal MRI for this kind of classification task. For instance authors of study (Bouts et al. 2018) used both structural and functional MRI (T1 weighted, Diffusion tensor, and resting state fMRI) and showed much higher performances (best AUC of 0.922) for classifying dementia-type differentiation vs normal brain. The model employed used a linear model (elastic net regularized regression). However this paper used on a small dataset (37 patients with probable AD, and 28 patients with behavioral variant frontotemporal dementia, vs 35 control subjects).

### About the limits of CNN

One limit of CNN resides in their lack of “common sense”, ie this algorithm can do misclassification that a human would never do. For instance, high confidence predictions were attributed to unrecognizable images (Nguyen, Yosinski, et Clune 2014). This idea was showed in a different way : adding a small “adversarial patch” (a small circle of unrecognizable image) (Brown et al. 2017) changed predictions. Similar results were obtained adding non perceptible variations in image (Szegedy et al. 2013).

In our results, we can link this concept to the fact that better results were observed with random split. The models captured the informations related to hospitals and used it for predicting age.

### Go beyond CNN : futur promises

Auto-Machine learning (Auto ML) is a promising evolution of machine learning. Today a lot of time consumed by data scientists consists in empirical fine-tuning of hyperparameters, or finding relevant digital architecture for a given problem to solve. An automatisation of theses aspects could accelerate the ML process. For instance, a neural network could be used… for discovering neural network architectures (for a recent review, see (Elsken, Metzen, et Hutter 2018)).

Capsule networks (Sabour, Frosst, et Hinton 2017) showed state-of-the-art performance on classical benchmark in Computer Vision classification task (MNIST). Contrary to CNN, its architecture takes into account spatial correlation in image (for instance, on a face detection task, nose is classified as a nose because capsule networks considered spatial information about where nose place in a human face, (classically found… between and below the eyes). Capsules network are designed for considering such spatial information. The promise of this property resides in relatively lower amount of data need for similar than CNN’s performance.

Deep reinforcement learning is a method inspired by dopaminergic reward system in brain, an recently outperformed alphago champion (Silver et al. 2016). This could be an interesting technique, but the main limit results in the high number of training examples required for efficient training. We can cite a paper focused on Lung nodule detection (Ali et al. 2018), as a proof of concept, but without actionable results yet (overall accuracy on test set of 64.4% (sensitivity 58.9%, specificity 55.3%, PPV 54.2%, and NPV 60.0%)).

To conclude, CNN showed powerful performances on computer vision task, and its applications seem promising for medical imaging. However, this models has to be used with caution, in particular because of their high sensitivity to bias present in images itselves. Collaborative work with machine learning expert and physicians seems to be an efficient way of doing proper science in order to find accurate and explainable biomarker.

